# Earliest-known intentionally deformed human cranial fossil from Asia and the initiation of hereditary hierarchy in the early Holocene

**DOI:** 10.1101/530907

**Authors:** Xijun Ni, Qiang Li, Thomas A. Stidham, Yangheshan Yang, Qiang Ji, Changzhu Jin, Khizar Samiullah

## Abstract

Hereditary hierarchy is one of the major features of complex societies. Without a written record, prehistoric evidence for hereditary hierarchy is rare. Intentional cranial deformation (ICD) is a cross-generational cultural practice that embodies social identity and culture beliefs in adults through the behavior of altering infant head shape. Therefore, ICD is usually regarded as an archeological clue for the occurrence of hereditary hierarchy. With a calibrated radiocarbon age of 11245-11200 years BP, a fossil skull of an adult male displaying ICD discovered in Northeastern China is among the oldest-known ICD practices in the world. Along with the other earliest global occurrences of ICD, this discovery points to the early initiation of complex societies among the non-agricultural local societies in Northeastern Asia in the early Holocene. A population increase among previously more isolated terminal Pleistocene/early Holocene hunter-gatherer groups likely increased their interactions, possibly fueling the formation of the first complex societies.

## Introduction

Human societies are thought to have become more complex since the beginning of the Holocene (about 11 ka) (1-3) after the end of the last major glacial interval. Hereditary hierarchy is generally regarded as one of the essential characteristics of complex societies. The origins of hereditary inequality, or the replacement of egalitarian societies with ascribed hereditary hierarchy, represents the major transition from a lower to a higher level of social complexity (2). Without written records, prehistoric evidence for this crucial transition in human cultural evolution is rare. However, intentional cranial deformation (ICD), also known as artificial cranial modification, generally is regarded as a definitive archeological signal for hereditary rank (4, 2, 5).

The human head is widely regarded as the carrier of the soul, identity, personhood, ancestry, and ethnicity (6). Past and present societies and cultures have treated the heads of living and deceased individuals in very diverse ways, including their decoration, deformation, trephination, and even decapitation (6). Within that diversity, ICD is an intriguing cultural practice that is a deliberate, significant, and permanent modification to the shape of the human skull, produced by compressing an infant’s head with hands, binding the head with hard, flat surfaces, or by tightly wrapping the head in cloth. ICD is a ritualized and cross-generational cultural practice that results in an easily recognizable flat, elongated, or a conical vault shape in adulthood. The practice of ICD and its associated meme are regarded as a significant way to symbolize social identity and embody cultural beliefs, with the results signifying group affiliation, or demonstrating social status (7).

ICD has a rich historic and prehistoric record in cultures around the world (8-12, 7) that probably continues today (13). Although the practice of ICD has been suspected in the Neanderthals (14) and a latest Pleistocene human from the Upper Cave of the Zhoukoudian locality in China (15), these suggestions have been seriously challenged (at least they were not intentinally deformed) (16, 17, 7). The earliest known preserved skulls that undoubtedly exhibit ICD are from the early Holocene (about 10000 – 12000 years ago) in the central Murray River region of southeastern Australia and the Proto-Neolithic deposits in Shanidar Cave in Iraq (18-20) (see Supplementary Information).

## Results

The new fossil clearly exhibiting ICD (Songhuajiang Man I, IVPP PA1683, Figure 1) reported here was collected at an underwater sand mining site in the Songhuajiang River near Harbin, Heilongjiang Province, Northeastern China. While the sedimentology and particular fossiliferous layer where the partial cranium was discovered are unknown, direct AMS radiocarbon dating of the cranium reveals a calibrated date of 11245-11200 years BP (Supplementary Information). That age places the Songhuajiang ICD record roughly as old as the previously-oldest known ICD records from Australia (calibrated AMS ^14^C date of 11440 ± 160 years BP, the most securely dated age) (18)and older than the records from the Middle East (calibrated ^14^C date of 10600 ± 300 years BP) (19, 20).

**Figure 1.**
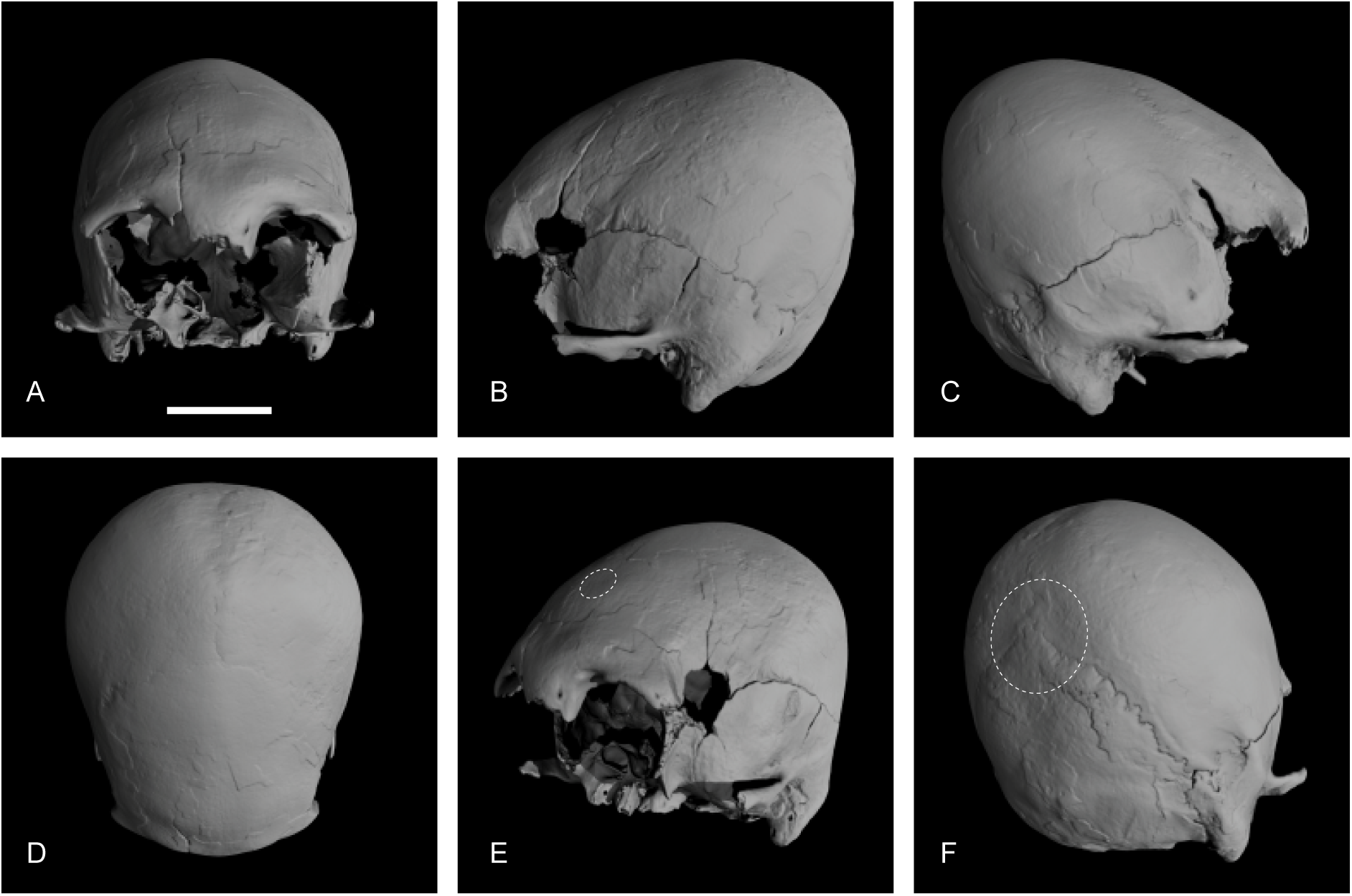
Songhuajiang Man I intentionally deformed cranium fossil (IVPP PA1683). A, Anterior view; B, Left lateral view; C, Right lateral view; D, Superior view; E, Anterior-left lateral view; F, Posterior-right lateral view. White dashed circles in E and F indicate the flat areas that likely were secured against a hard surface during infancy/early childhood. White scale bar indicates 5 cm.

The Songhuajiang ICD cranium is lightly built, showing modern aspects that resemble the Proto-Neolithic crania from Shanidar Cave and Jericho (19, 20), but differing from the heavily built crania from southeastern Australia (18). The facial cranial part of the skull is missing, but the neurocranial portion of the skull is nearly complete. Only the sphenoid and the condyles of the occipital are damaged. The individual has a prominent supraorbital ridge, blunt supraorbital margin, strong temporal line, and a long and large mastoid process, suggesting that the cranium probably belongs to a male (21, 22). The preserved cranial sutures show early stages of fusion. The coronal and sagittal sutures exhibit minimal to marked degree of closure, while the other sutures, such as lambda and sphenotemporal, are almost unfused. Based on the composite scores reflecting the degree of the ectocranial suture closure (23), the age-at-death is estimated as young adult (20-34 years). The assessment of its ethnicity is difficult, given the missing facial cranial region and its antiquity, but the moderately wide interorbital space, large mastoid process, shallow infraglabellar notch, and rounded and sloping supralateral margin of the orbits commonly are present in Asian people, particularly in the modern Chinese population (24-26). Despite the strong deformation of the parietal region, the skull shows a clear obelionic depression near the parietal foramen. The frequency of occurrence of this feature is very high in the modern Chinese population (24, 25). Overall, these non-metric cranial characteristics mentioned above are more consistent with the morphology of a young male Asian individual.

The oldest-known ICD records from Australia exhibit a very mild change to cranial shape, likely resulting from pressing hands onto the newborn’s forehead (27, 18). The oldest-known, undoubted ICD records from the Middle East were generated by circumferential restriction to skull growth by wrapping the infant’s head with one or more pieces of cloth. By contrast, the Songhuajiang Man I exhibits typical tabular deformation, which clearly is a more complicated and sophisticated ICD methodology than the manual molding or annular type of deformation. This version of ICD results in a cranial vault, as seen in the Songhuajiang Man I fossil, that has a core-like shape, with a very flat and superior-posteriorly sloping forehead, and a nearly vertical, flat occiput. The forehead does not exhibit a circular depression that would have formed as the result of the restriction by bandages (as in annular deformation). The squama of the frontal bone is flat with a very low frontal eminence. The frontal curvature index (a measure of the flatness calculated as the ratio of the subtense height relative to the chord length) is 14.1, much lower than in undeformed skulls (18) (Supplementary Information). There is a small flat area between the two frontal eminences, and displaced slightly to the left side (Figure 1e). This flat region probably is the result of binding a hard flat plate or board (possibly a so-called anterior pillow (11)) on to the frontal region for a prolonged time during the growth of the immature skull. The parietal is anteroposteriorly shortened, but superior-posteriorly elongated. The parietal curvature index is 27.6, much higher than that of undeformed skulls (18). As in other tabularly deformed crania, the parietal eminence bulges significantly laterally and has a large curvature. In superior view, the posterior part of the skull is much wider than the anterior part, and clearly differs from the cylindrically shaped cranium produced by annular deformation. There is a disk-like flat region around the lambda (Figure 1f), which must be the result of the area being secured against a hard flat surface during infancy/early childhood. The squama of the occipital region also is flat. The occipital curvature index is 18.6, much lower than in undeformed skulls (18). The external occipital protuberance and external occipital crest are weak.

Cranial deformation affects the frequency of wormian bones (suture ossicles) (28-31), and also alters the endocranial structures such as brain shape and blood vessel patterns (32-34). The Songhuajiang Man I has a small piece of wormian bone along the parietal-mastoid suture, and a tiny piece on the occipital-mastoid suture on the right side of the skull. Although the wormian bones on the lambda suture commonly are present in modern African and European populations, they have a lower frequency in the extant Asian population (35). The virtual endocast of the Songhuajiang Man I (Figure 2) is anterior-posteriorly shorter than an undeformed endocast. The frontal and occipital regions are flatter, while the parietal region is more superiorly projecting. In lateral and posterior view, the occipital transverse sulcus and lunate sulcus are broad and deep. In undeformed endocranial casts, these two sulci are usually invisible or quite shallow (36). On the left side, the anterior branch of the middle meningeal vessel is deep, thick, and has many crenulated terminal branches. The posterior branch of the middle meningeal vessel is shallower, thinner and has fewer branches. As it is commonly seen in other deformed crania, the posterior part of the posterior branch angles superiorly and extends to the top of the parietal lobe. On the right side, the anterior branch of the middle meningeal vessel is as deep and complicated as that on the left side. The entire posterior branch alters its direction by extending posterior-superiorly, almost parallel to the anterior branch. The anterior and posterior branches are similarly deep, but the posterior branch has fewer branches. In the undeformed endocast, the anterior and posterior branches of the middle meningeal vessel are equally thick, moderately deep, lack the convoluted terminal branches, and do not exhibit the altered direction (32-34). The impression of the sagittal sinus of the Songhuajiang Man I is quite deep and does not display compression or flattening, different from most other deformed crania. The drainage predominantly goes to the right transverse sinus. In undeformed crania, the sagittal sinus usually drains to the right transverse sinus, while in deformed crania, it has a higher frequency of travelling to the left side (32, 34).

**Figure 2.**
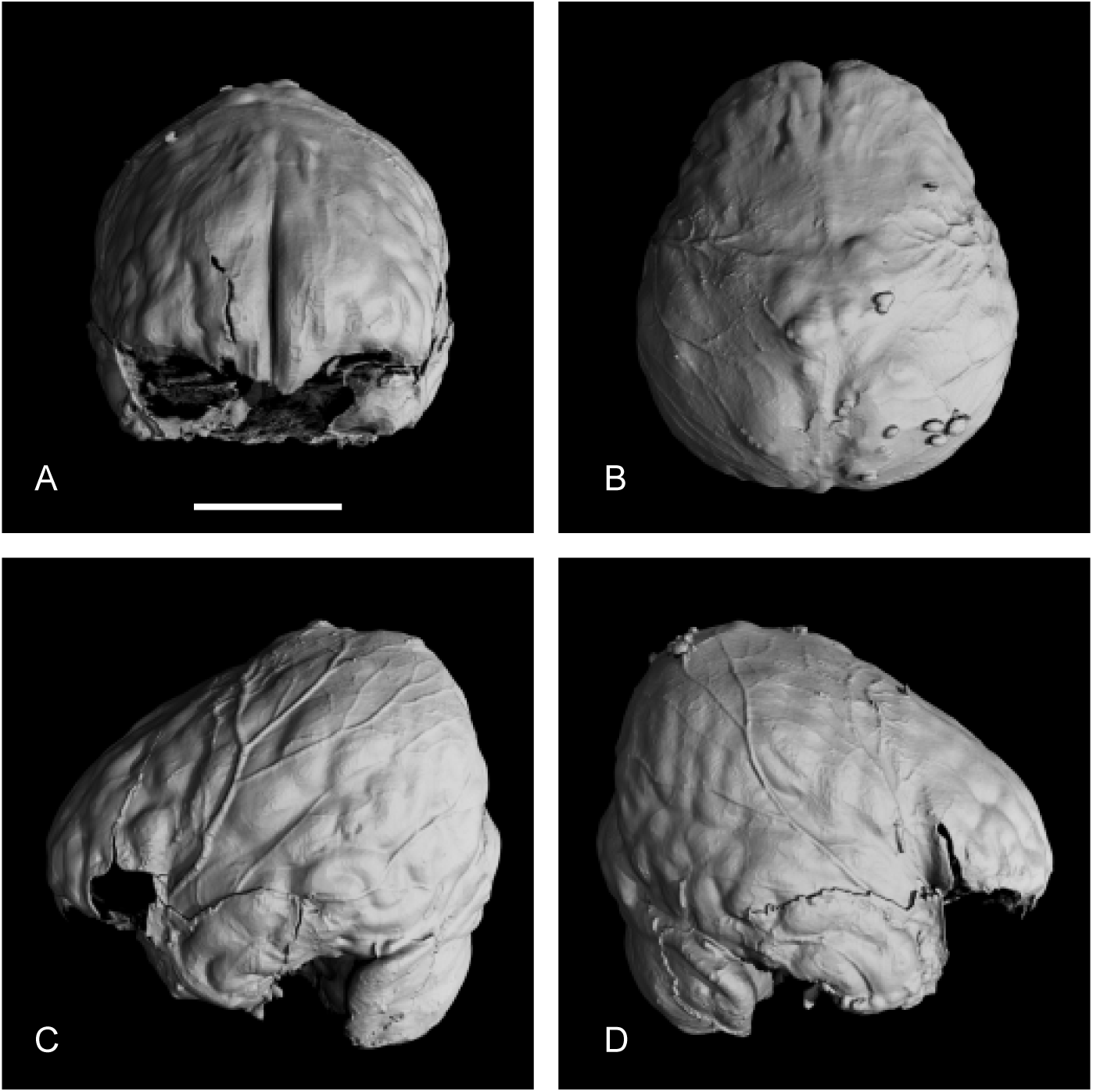
Virtual endocast of the Songhuajiang Man I (IVPP PA1683). A, Anterior view; B, Superior view; C, Left lateral view; D, Right lateral view. White scale bar indicates 5 cm.

It is widely assumed that the variation among individual people in the isotopic composition (δ^13^C-δ^15^N) of their bone collagen reflects differences in their diets (37, 38). The δ^13^C of Songhuajiang Man I is −20.8 ‰, and the δ^15^N is 11.4 ‰. The low carbon isotope value indicates that this man had a diet mainly consisting of C_3_ foods. Nitrogen stable isotope ratios are low among terrestrial herbivores, but high in fishes (39, 37). The relatively high nitrogen isotope value in the bone collagen of Songhuajiang Man I not only suggests his dietary protein originated from high trophic levels, but also indicates that the animal protein probably derived from sources in lakes and rivers. The isotopic composition of δ^13^C and δ^15^N in Songhuajiang Man I is very close to the values found among the early and middle Neolithic people from the Angara River and Upper Lena River in Siberia (39), who were hunter-fisher-gatherers.

In Australia, the Middle East, and now the Songhuajiang area, ICD crania make up only a small proportion of the known human specimens (Supplementary Information). That low frequency likely is the result of only select individuals in these populations (maybe the elite children) being involved in the practice and application of ICD, suggesting that ICD was a very selective behavior, and probably also a ritual action, before it became a meme widely distributed across different time periods and cultures. This selectivity is highly suggestive of the presence of social and/or cultural differentiation within the populations or cultures. Additionally, these earliest ICD records show that different geographically distant, early cultures around the world spread this cultural practice from one generation to the next outside of an agricultural setting. The association of ICD with agricultural practice appears to have occurred later in the Holocene.

## Discussion

Various ritualized and costly actions, such as firewalking, ritual scarification, and subincision, are theorized as credibility enhancing displays that could promote group solidarity and intragroup cooperation, and they reflect a deep level of commitment to group ideologies and religious beliefs (40). The practice of ICD is such a ritualized and costly action that results in highly visible and permanent life-long body modification that even extends beyond an individual’s lifespan (41, 42, 7). Differing from other typical body treatments, ICD is performed by the parental generation and applied to the descendant generation (a cross-generational action). Through this practice, a highly visible and permanent icon for specific social identity and cultural beliefs is embodied from one generation to the next generation. The continuation of this cultural tradition requires much forethought and planning to achieve the desired result in adults of the next generation. Individuals with their appearance, as the result of ICD, obtained an ascribed status that was enhanced by an easily recognizable, highly visible, and immutable physical trait. The practice symbolized in ICD is not only accepted and understood by the clan/group who performed this activity on their descendants, it also should be understood and recognized by other groups who have interaction with the ICD expressing groups, including those who do not perform this practice.

As in Australia and the Middle East, the practice of ICD in the Songhuajiang area occurred in hunter-gatherer populations. The isotopic composition of δ^13^C-δ^15^N of the bone collagen of Songhuajiang Man I indicates that the man had a diet mainly consisting of C_3_ foods and ingested protein that originated from sources in lakes and rivers. It is generally believed that hunter-gatherers tend to have an egalitarian social ethos. However, by reinforcing the ascribed status of a social entity via such a sophisticated and ritualized way as the practice of ICD, the people in these areas signified their intragroup affiliation and delimited intergroup social boundaries. Multiple clans, with and without the practice of ICD, must have had frequent interactions, and probably formed a kind of larger tribe or community. Relatively stable, but complex and likely socially stratified tribes or communities formed by different hunter-gatherer clans likely were present in the Middle East, Australia, and East Asia during the Pleistocene-Holocene transition.

The natural and social factors that led to the practice of ICD, and the intertwined enhanced ascribed social status resulting from this practice are critical for understanding the early evolution of human social complexity. Even though it is widely suggested that the development of intensive resource use, such as the shifts from foraging to agriculture, is relevant to the (origin and) evolution of social complexity, recent anthropological research suggests that intensification of resource use or acquisition and sociopolitical hierarchy are broadly reciprocal, probably as a feedback loop that also may have involved population growth (3).

The terminal Pleistocene-early Holocene transition is marked by a major period of significant climate change, forming an ecological threshold for humans. A penecontemporaneous dramatic increase in the human population occurred independently in different parts of the world, presumably when humans adapted to a more sedentary lifestyle (43, 44), and this increase is widely considered as the result of the agricultural revolution. However, sudden increases in human and artifact remains in areas without evidence of agriculture, such as Siberia, Northeast China, and Australia, suggests that the spurt of demographic growth also was present at the same time in areas with predominantly hunter-gatherer groups (45, 46). The population increase among these hunter-gatherer populations may have increased the home ranges of various clans and propelled their dispersal. This change consequently increased the likelihood of interactions and communication among previously more isolated hunter-gatherer groups. The practice of ICD for enhancing social identity in these areas likely reflects their social complexity, as well as inequality or disparity among individuals (i.e. stratification). It is generally accepted that social inequality and complexity are correlated with the degree of social stratification (47). A socially stratified group or tribe is more organized and efficient in a variety of endeavors including hunting, gathering, and agricultural production. The cross-generational nature of ICD (and its associated meme) actually may have functioned to reinforce that social stratification or to increase cultural cohesion continuing the social complexity or hierarchy over time, in particular during the transition to more sedentary and agriculturally oriented societies in the early Holocene.

Given the current archeological record, the oldest-known evidence for the practice of ICD is spread over less than a thousand years across a very large geographic area (Middle East, East Asia, and Australia, Figure 3) close in time to the Pleistocene-Holocene transition (∼11.7 ka) (18-20). Despite that narrow temporal window, the methods for deforming a neonate’s cranium differed among those three populations. While it cannot be totally rejected that the people from about 11,000 years ago in Australia, Northeast Asia, and the Middle East shared a deep social belief or identity related to ICD expression, it is more likely that the people from these three widely separated areas faced a similar incentive that drove them to make similar cultural responses. However, the later prevalence of ICD among native populations and cultures of the Americas may have a different interpretation. Genetic studies have revealed unquestionable Asian origins of the natives in the New World, and the archaeological records in Siberia and Beringia suggest that the initial dispersal of modern humans to North America probably occurred after 15 ka (48, 49). Given those close genetic and cultural connections, the widespread native practice of ICD in the New World may derive from a continuation of this potentially ancestral cross-generational meme in northeastern Asia.

**Figure 3.**
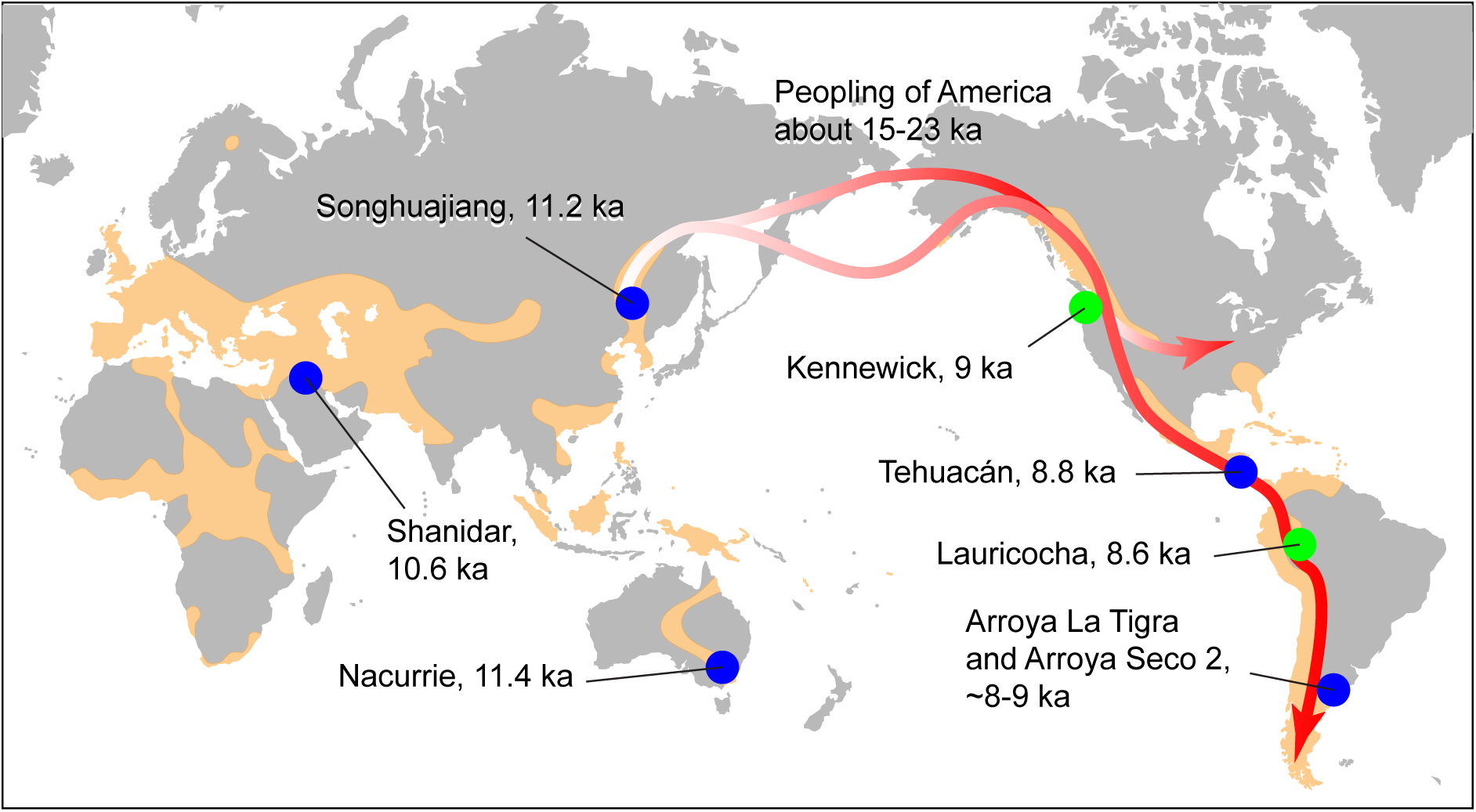
Worldwide distribution of the practice of intentional cranial deformation (ICD) in archeological records. The areas in pale orange indicate the distribution. Blue dots indicate sites with undoubted ICD. The green dots indicate places with dolichocephalic crania, which are elongated heads that may not be ICD. Data from references (18, 8-11, 19, 50, 12, 20, 7). The date for peopling of Americas is inferred from analyses of genomic data from reference (49). The global map is modified from https://upload.wikimedia.org/wikipedia/commons/c/c2/Blank_Map_Pacific_World.svg (under the Creative Commons Share Alike license: https://creativecommons.org/licenses/by-sa/3.0/deed.en).

## Material and Methods

Supplementary Information provides full information about CT scanning, radiocarbon dating, and trophic position (stable isotope) analysis.

## Acknowledgements

This project has been supported by the Strategic Priority Research Program of Chinese Academy of Sciences (CAS XDB26030300, XDA19050100, XDA19050102), the National Natural Science Foundation of China (41472025, 41625005), and the External Cooperation Program of BIC (132311KYSB20160008). We are grateful to Yemao Hou for CT scanning. Drs. Tao Deng, Wu Liu provided instructive suggestions.

## Competing Interests

The authors declare no competing interests as defined by eLife, or other interests that might be perceived to influence the results and/or discussion reported in this paper.

