## Supplementary Information for "Earliest-known intentionally deformed human cranial fossil from Asia and the initiation of hereditary hierarchy in the early Holocene"

**Title:**

Intentional cranial deformation; complex society; cross-generational cultural practice; hereditary hierarchy

1. **Oldest-known Intentional Cranial Deformation Records**

The cultural practice of intentional cranial deformation (ICD) has a very large geographic distribution, and a very long documented history extending into the late Pleistocene. It is present on all continents at various periods in human history(1-6), and probably continues today (in Vanuatu(7)). The oldest specimens suspected of having this special body treatment are the 45 ka Shanidar 1 and Shanidar 5 Neanderthal skulls (8). Although this hypothesis is still widely cited, the original author subsequently pointed out in one of his coauthored papers that the evidence of ICD in the Shanidar 5 is not present (9), while the presence in the Shanidar 1 skull also is questionable. A recent quantitative analysis failed to support the Shanidar 1 skull as exhibiting ICD (10). A skull from the Upper Cave of the Zhoukoudian locality in China (Upper Cave 102) is another widely cited ICD record. A shallow depression anterior to the coronal suture was interpreted as the result of intentional banding (11). The preserved cast of the specimen in IVPP shows clear postmortem bilateral compression. Other than the shallow depression, no other typical ICD features appear to be present in that specimen. Even though the skull may have been deformed during life, there is no reason to believe that it was intentionally modified (6). Along with many other Zhoukoudian *Homo* specimens, Upper Cave 102 was lost during World War II, and it is not possible to test whether the depression anterior to the coronal suture was the result of postmortem deformation. The age of that specimen also is in question. Radiocarbon dating on the samples of bones taken from the sediments in stratigraphic order indicates an age spanning between 34 and 24 ka, and 29 - 24 ka for the cultural layers (12). However, the hominin remains were suggested to be part of the intentional burials, and thus should be younger than the naturally preserved animal bones from the same layer (13).

Excavations at the Nacurrie, Coobool Creek, and Kow Swamp burials in the central Murray River region of southeastern Australia produced some interesting evidence of ICD. A few skulls, such as Kow Swamp 5, Coobool Creek 65, and Nacurrie 1, show a combination of features including a flat frontal, minimum frontal breadth posteriorly positioned, distinct pre-bregmatic eminence, and low position of the maximum cranial breadth. These features are thought to be suggestive for the regional evolutionary sequence from Indonesian *Homo erectus* to the Australian *H. sapiens* (14, 15). However, most subsequent researches on the larger and better preserved samples from the Nacurrie, Coobool Creek, and Kow Swamp burials suggest that the shape of the peculiar cranial vault shape (as in Kow Swamp 5 and other skulls with a flat frontal from these localities) is the result of ICD, rather than normal growth processes (16-20, 10, 21). These human remains from the central Murray River region were dated as terminal Pleistocene to early Holocene (19). Nacurrie 1 is the most securely dated burial, given that collagen extracted from a postcranial bone fragment of this individual has a calibrated AMS ^14^C date of 11440 ± 160 years BP (20).

Besides the questionable ICD skull from Zhoukoudian, another possible earliest-known ICD record from Eurasia is from the Arene Candide cave in Italy. Excavations at this site unearthed about twenty skeletons buried at the bottom of a layer that contains a Late Epigravettian industry and has a calibrated ^14^C date between 10910 ± 90 years BP and 11750 ± 95 years BP. One fragmentary skull from this cave seems have a particularly elongated neurocranium, and was hypothesized to have been subjected to oblique circumferential banding (22). However, a large part of the vault is missing. A later study revealed no sign of the effect of oblique bandaging, and suggested that the skull deformation probably was caused by an accident (23).

A slightly younger specimen from the Proto-Neolithic deposits (Layer B1, with a calibrated ^14^C date of 10600 ± 300 years BP) in Shanidar Cave in Iraq is generally accepted as the earliest Eurasian record of ICD (24, 25). ICD practice also is known from the Pre-Pottery-Neolithic A and B (10300-7300 BP, calibrated ^14^C date) of the Levant. In Jericho, the ICD practice was discovered associated with mortuary practices, such as skull removal, decoration, and caching (26). Fletcher et al. ((26)) examined a plastered skull (D113, 10,100–9250 calibrated ^14^C years BP) unearthed from the Kathleen Kenyon at Neolithic Jericho, and revealed that the skull exhibits ICD (26). They suggested that the post-mortem treatment of an ICD skull is a ritual practice that was of significance during life, not just after death (26).

The ICD habit has a long and probably continuous history in Northeast Asia. Previously oldest-known ICD human in East Asia is from the Qingshantou Locality in Jilin Province (with a ^14^C date of 9860 ± 160 years BP) (27). The Qingshantou locality is very close geographically to the Harbin site (about 200 km southwest). A well-preserved skull discovered from the Djalai-Nor coalfield (28, 29) shows clear anteroposterior (fronto-occipital) compression. There are at least two fossil layers at the Djalai-Nor coalfield. The human fossils were discovered in the upper layer and were found with a pottery sherd (30). That deformed skull currently is missing, and direct dating of the specimens is impossible. However, it is widely accepted that the human fossil found together with cultural remains is younger than ~10 ka (31-33). Between five and six thousand years ago, the occurrence of ICD became much more prevalent in North Asia. ICD individuals are known from the Boisman 2 Neolithic cemetery (ca. 5800–5400 ^14^C BP) in the far East of Russia (34). During the Dawenkou Culture period (about 6300-4500 years ago) in China, human cranial remains usually exhibit a flat occipital region, but do not have a flat frontal region (35-38). It is widely believed that the flat occipital in Dawenkou people is the result of placing babies with their backs against a hard, flat surface.

The archeological records of the ICD practice in the Americas are a few thousands of years later than those from Asia and Australia. The oldest-known record is from the Central Highland Basin of Tehuacan in Mexico, with a date about 8800 years BP (6). The cranium from the Arroyo La Tigra in Argentina shows some signs of the practice of ICD (39) (but see ref. (40)). The calibrated radio-carbon age of the Arroyo La Tigra is 8180-7960 BP (39). With a slightly younger age (uncalibrated radiocarbon date, 7615±90 BP), the Arroyo Seco 2 archeological site show quite high proportion of undoubted ICD human remains (41). A partial cranium discovered from a layer about 8550 years old in the Lauricocha cave in the Central Andean highlands in Peru (42) and the skull of Kennewich man recovered near Kennewick, Washington (radiocarbon dated to 8340–9200 calibrated years BP) (43, 44) are described as the “dolichocephalic” crania. This specific cranial form is usually regarded as the key feature that distinguishes Paleoamericans from the modern Native Americans (42, 45). They were not considered as the ICD records, however, both of the crania show a shallow depression posterior to the bregma, which is a character usually considered as the result of the practice of ICD.

2. **Radiocarbon dating and trophic position**

A piece of bone was removed from the left side of the occipital. The sample was analyzed at the Beta Analytic Inc. in Miami, Florida. Results are ISO/IEC-17025:2005 accredited. Bone collagen was extracted by using alkali and solvent extraction. Accelerated Mass Spectrometry (AMS) measurements were made on one of 4 in-house NEC SSAMS accelerator mass spectrometers. The reported age is the "Conventional Radiocarbon Age", corrected for isotopic fraction using the 𝛿 ^13^C. Age is reported as radiocarbon years before present (BP). "Present" is 1950 AD. By international convention, the modern reference standard was 95% the ^14^C signature of NBS SRM-4990C (oxalic acid) and calculated using the Libby ^14^C half life (5568 years). Quoted error on the BP date is 1 sigma (1 relative standard deviation with 68% probability) of counting error (only) on the combined measurements of sample, background and modern reference standards. Total error at Beta (counting + laboratory) is known to be well within +/- 2 sigma. 𝛿 ^13^C values are reported in parts per thousand (per mil) relative to PDB-1 measured on a Thermo Delta Plus IRMS. Typical 𝛿 ^13^C error is +/- 0.3 o/oo. Percent modern carbon (pMC) and 𝛿 ^14^C are not absolute. They equate to the Conventional Radiocarbon Age. Calendar calibrated results were calculated the material appropriate 2013 database (INTCAL13, MARINE13 or SHCAL13). The trophic position of the Songhuajiang Man I was evaluated by stable isotope analysis. The reported 𝛿 ^13^C values were measured separately in an IRMS (isotope ratio mass spectrometer). They are NOT the AMS 𝛿 ^13^C which would include fractionation effects from natural, chemistry and AMS induced sources.

The conventional radiocarbon age of the Songhuajiang Man I skull is 9810 ± 30 BP. Calibrated result (95% probability) is 9295-9250 BC, or 11245-11200 BP. The 𝛿 ^13^C of Songhuajiang Man I is -20.8 ‰, and the 𝛿 ^15^N is 11.4 ‰.

3. **CT-scanning and measurements**

The Songhuajiang Man I ICD cranium fossil (IVPP PA1683) were CT-scanned using the 450kv High Resolution CT facility at the Key Laboratory of Vertebrate Evolution and Human Origins, Chinese Academy of Sciences. Segmentation and 3D virtual reconstructions of the skull and the endocast were done by following the standard procedure introduced by Ni et al. (46). Measurements (Table S1) were taken based on the virtual reconstruction in VG Studio Max 3.1.

Table S1. Linear dimensions and subtenses in the Songhuajiang Man I. Measure definition as in reference (20)

| Variables | Measurements |
| --- | --- |
| Frontal chord (n-b) | 126.07 mm |
| Frontal sub h | 17.78 mm |
| Frontal curvature index | 14.1 |
| Frontal angle | 19.84 |
| Parietal chord | 96.28 mm |
| Parietal sub h | 26.61 mm |
| Parietal curvature index | 27.6 |
| Parietal angle | 25.6 |
| Occipital chord | 110.59 mm |
| Occipital sub h | 20.55 mm |
| Occipital curvature index | 18.6 |
| Occipital angle | 18.09 |
| Basion-bregma | 154.45 mm |
| Max biparietal br | 141.68 mm |
| Min frontal br | 92.5 mm |
| Nasion-opisthocranion | 169.84 mm |

4. **Reference**

1.	Dingwall EJ (1931) *Artificial Cranial Deformation, a Contribution of the Study of Ethnic Mutilations* (John Bale, Sons & Danielsson, LTD., London) p 313.

2.	Enchev Y, Nedelkov G, Atanassova-Timeva N, & Jordanov J (2010) Paleoneurosurgical aspects of Proto-Bulgarian artificial skull deformations. 29(6),E3:1-7. *Neurosurgical Focus* 29(6):E3:1-7.

3.	Jung H & Woo EJ (2017) Artificial deformation versus normal variation: re-examination of artificially deformed crania in ancient Korean populations. *Anthropological Science* 125(1):3-7.

4.	Kiszely I (1978) *The origins of artificial cranial formation in Eurasia from the sixth millennium B.C. to the seventh Century A.D.* (B.A.R., Oxford) p 76.

5.	Ricci F*, et al.* (2008) Evidence of artificial cranial deformation from the later prehistory of the Acacus Mts. (southwestern Libya, Central Sahara). *International Journal of Osteoarchaeology* 18(4):372-391.

6.	Tiesler V (2014) *The bioarchaeology of artificial cranial modifications* (Springer, New York).

7.	Childress DH & Foerster B (2012) *The enigma of cranial deformation* (Adventures Unlimited Press, Kempton).

25.	Solecki RS, Solecki RL, & Agelarakis AP (2004) *The Proto-Neolithic Cemetery in Shanidar Cave* (Texas A&M University Press, College Station).
